# Improving the immunogenicity of *E. coli* FimH via multivalent display on I53-50 nanoparticles

**DOI:** 10.64898/2026.03.05.709951

**Authors:** Rebecca S. Cole, Natalie C. Silmon de Monerri, Jacqueline Lypowy, Christopher Ponce, Cara Kobylarz, Lily Liu, Zein Kasbo, Elizabeth Kepl, Tara Ciolino, Art Illenberger, Leslie Gallardo, Annalena Laporte, Danielle Baranova, Rashmi Ravichandran, Laurent O. Chorro, Robert G.K. Donald, Raphael Simon, Neil P. King

**Affiliations:** Institute for Protein Design, University of Washington, Seattle, WA, USA; Department of Biochemistry, University of Washington, Seattle, WA, USA; Vaccine Research and Development, Pfizer Inc, Pearl River, NY, USA

## Abstract

Urinary tract infections, caused primarily by uropathogenic *E. coli*, are a significant public health burden, affecting approximately 50% of women worldwide. The adhesin FimH is responsible for host receptor binding and is therefore a promising vaccine candidate, but prior studies showed that recombinant monomeric FimH is poorly immunogenic. Here we displayed FimH antigens on the two-component protein nanoparticle I53-50 to generate nanoparticle immunogens that elicit robust levels of receptor-blocking antibodies in mice and non-human primates. We produced nanoparticle immunogens displaying either the FimH lectin domain or a recently reported conformationally stabilized antigen, FimH-DSG, comprising both the lectin and pilin domains. When formulated on aluminum hydroxide, both nanoparticle immunogens elicited similar levels of receptor-blocking activity as a ten-fold higher dose of monomeric FimH-DSG formulated with a potent adjuvant. The improved manufacturability of the stabilized antigen, combined with the ability of nanoparticle display to obviate the need for complex adjuvants, provides important preclinical data for FimH-based vaccines intended to prevent urinary tract infections. More broadly, our data extend the applicability of the I53-50 nanoparticle platform, which to date has been mainly used for displaying viral and protozoan antigens, to bacterial indications.

## Introduction

Urinary tract infections (UTI) are the most common outpatient infection, affecting an estimated 150 million people annually^1,2^. While antibiotic treatments are available and effective, recurrent infections are common, particularly in the elderly^1^. Around 95% of infections are caused by uropathogenic *E. coli* (UPEC), which often colonizes the gut and can ascend the urinary tract via the urethra to establish a UTI. Colonization of the urinary tract is facilitated by long proteinaceous extensions called pili, of which Type 1 pili are the best studied. Type 1 pili are multimeric complexes comprising polymers of FimA subunits capped by FimF, FimG, and an adhesin on the tip called FimH, which binds to mannosylated uroplakins that cover the urinary tract epithelium^3^. Recombinant monomeric FimH has been explored as a vaccine target in several studies in mice and non-human primates, which showed that immunization is capable of eliciting receptor-blocking antibodies, although multiple doses and potent adjuvants were required^4–6^. The safety and immunogenicity of an adjuvanted FimCH protein complex has also been evaluated in a Phase I clinical trial among patients suffering from chronic UTIs. In this study, four doses of vaccine were required to induce anti-FimH immune responses and a measurable reduction in UTI incidence^7–9^. Collectively, these studies have identified FimH as a promising vaccine candidate, but that further optimization is needed to increase the antigen immunogenicity.

FimH consists of two domains: an adhesin or lectin domain (FimH_LD_) that mediates adhesion to bladder superficial umbrella cells by binding to mannosylated uroplakin Ia receptors on that coat the epithelial surface, and a pilin domain (FimH_PD_) that supports FimH_LD_ and connects it to the rest of the pilus via donor-strand complementation to the neighboring subunit, FimG^3,10^. FimH_LD_ can adopt low-affinity “open” and high-affinity “closed” conformations depending on the presence of ligand and the pilin domain, which acts as an allosteric regulator. FimH switches from low-affinity to high-affinity states with conformational changes that result from separation of the lectin and pilin domains as the FimH stretches due to mechanical force experienced during periods of high shear with urine flow^11^. In the absence of the pilin domain, FimH_LD_ spontaneously adopts the closed conformation. In constructs containing the pilin domain (including the FimG N-terminal donor strand), FimH_LD_ adopts the open conformation in the absence of ligand, but converts to the closed conformation when receptor is present^11^. The open conformation, which has a wider mannose-binding pocket, may be a better vaccine target as it is presumably the “prebound” conformation and could efficiently induce antibodies against the mannose binding site that prevent receptor binding. A recently described stabilized version of full-length FimH in which the lectin domain is locked in the open conformation, called FimH-DSG, can be expressed in high yield from mammalian cells, purified to homogeneity, and induces higher levels of inhibitory antibodies than FimH_LD_ (ref. ^12^). FimH-DSG therefore represents a promising stabilized FimH antigen that could be combined with complementary vaccine technologies to further improve its immunogenicity^13^.

Multivalent antigen display on self-assembling protein nanoparticles is a powerful strategy for enhancing the immunogenicity of subunit vaccines. Particulate immunogens have been shown to promote efficient lymphatic trafficking, potent B cell activation via multivalent receptor crosslinking, and improved germinal center formation^14–16^. This approach is particularly valuable for small, monomeric antigens like FimH, which often elicit suboptimal immune responses when administered in soluble form in the absence of coformulation with potent adjuvants. In previous work, we have shown that repetitive antigen display on computationally designed two-component I53-50 nanoparticles increases the magnitude, quality, and durability of vaccine-elicited antibody responses against a number of different viral and protozoan pathogens^14,17–19^. Furthermore, multiple I53-50-based immunogens have proven safe and immunogenic in clinical trials, and a vaccine comprising the I53-50 nanoparticle displaying the SARS-CoV-2 Spike receptor binding domain (RBD) has been licensed for use in multiple countries^20–22^. However, the utility of I53-50 in improving responses against bacterial antigens has only recently begun to be explored^23^.

Here we investigated the ability of I53-50 to improve the immunogenicity of FimH_LD_ and FimH-DSG in mice and non-human primates. We found that display on I53-50 significantly increased the magnitude of both binding and functional antibody responses relative to soluble FimH antigens, particularly after one or two doses of vaccine, establishing FimH-I53-50 nanoparticle immunogens as promising candidates for further development.

## Results

### Design, *in vitro* assembly, and characterization of FimH-bearing protein nanoparticle immunogens

We set out to produce I53-50–based nanoparticle immunogens displaying FimH and evaluate their ability to induce potent receptor-blocking antibody responses (**Fig. 1a**). To compare FimH_LD_ and FimH-DSG when multimerized, we generated constructs in which each antigen was genetically fused to the N terminus of I53-50A, the trimeric subunit of the I53-50 nanoparticle^24^, via a flexible linker (**Fig. 1b** and **Supplementary Table 1**). Both proteins were secreted from mammalian cells, and size exclusion chromatography (SEC) indicated that each formed monodisperse trimers as expected.

**Fig. 1.**
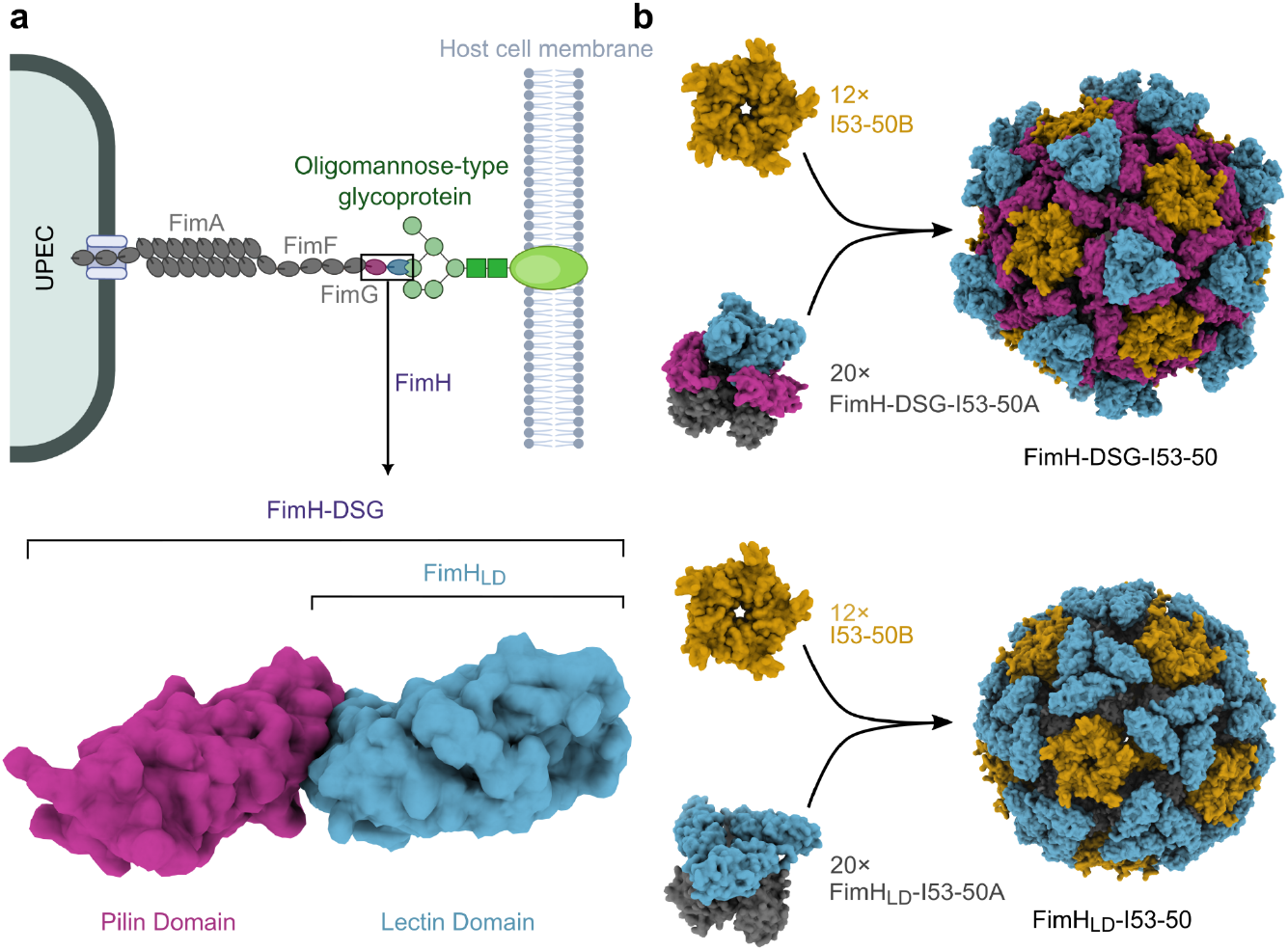
Design and assembly of FimH-bearing protein nanoparticles. **a** Structure of the type 1 pilus of UPEC in complex with an oligomannose-type glycoprotein (*top*), and close-up view of the molecular structure of the fimbrial adhesin FimH (*bottom*). **b** AlphaFold2 prediction of FimH-DSG-I53-50 and FimH_LD_-I53-50 nanoparticle immunogens, with pentameric I53-50B in orange and trimeric I53-50A in gray. The FimH_LD_ and FimH-DSG antigens are colored as in **a**. The predictions show the antigens flush with the surface of the nanoparticle, but in solution the GS linker is expected to render the antigens flexible relative to the nanoparticle scaffold. Protein models were rendered using ChimeraX and the schematic in panel **a** was partially created in BioRender^25,26^.

Following previous work, we mixed each antigen-bearing component with I53-50B, the pentameric component of the I53-50 nanoparticle, in a 1.1:1 molar ratio to drive assembly of nanoparticles displaying 60 copies of FimH_LD_ or FimH-DSG^27,28^ (**Fig. 1b**). Preparative SEC revealed predominant peaks at the expected elution volumes, suggesting successful *in vitro* assembly (**Fig. 2a**). SDS-PAGE confirmed the presence of both components in these peaks (**Fig. 2b**), and negative stain electron microscopy (nsEM) revealed fields of monodisperse particles with the known I53-50 size and morphology^19,24^ (**Fig. 2c**). Dynamic light scattering (DLS) before and after freeze-thaw confirmed the samples were homogenous and monodisperse, with observed diameters of 34 nm for FimH_LD_-I53-50 and 38 nm for imH-DSG-I53-50 that closely match the expected diameters (**Fig. 2d**). Finally, both nanoparticles bound mAb 926^29^ similarly, suggesting that each antigen remained conformationally intact after assembly, purification, and freeze-thaw (**Fig. 2e**). We note that the avidity inherent in a bivalent IgG binding to a multivalent nanoparticle immunogen prevented quantitative affinity determination.

**Fig. 2.**
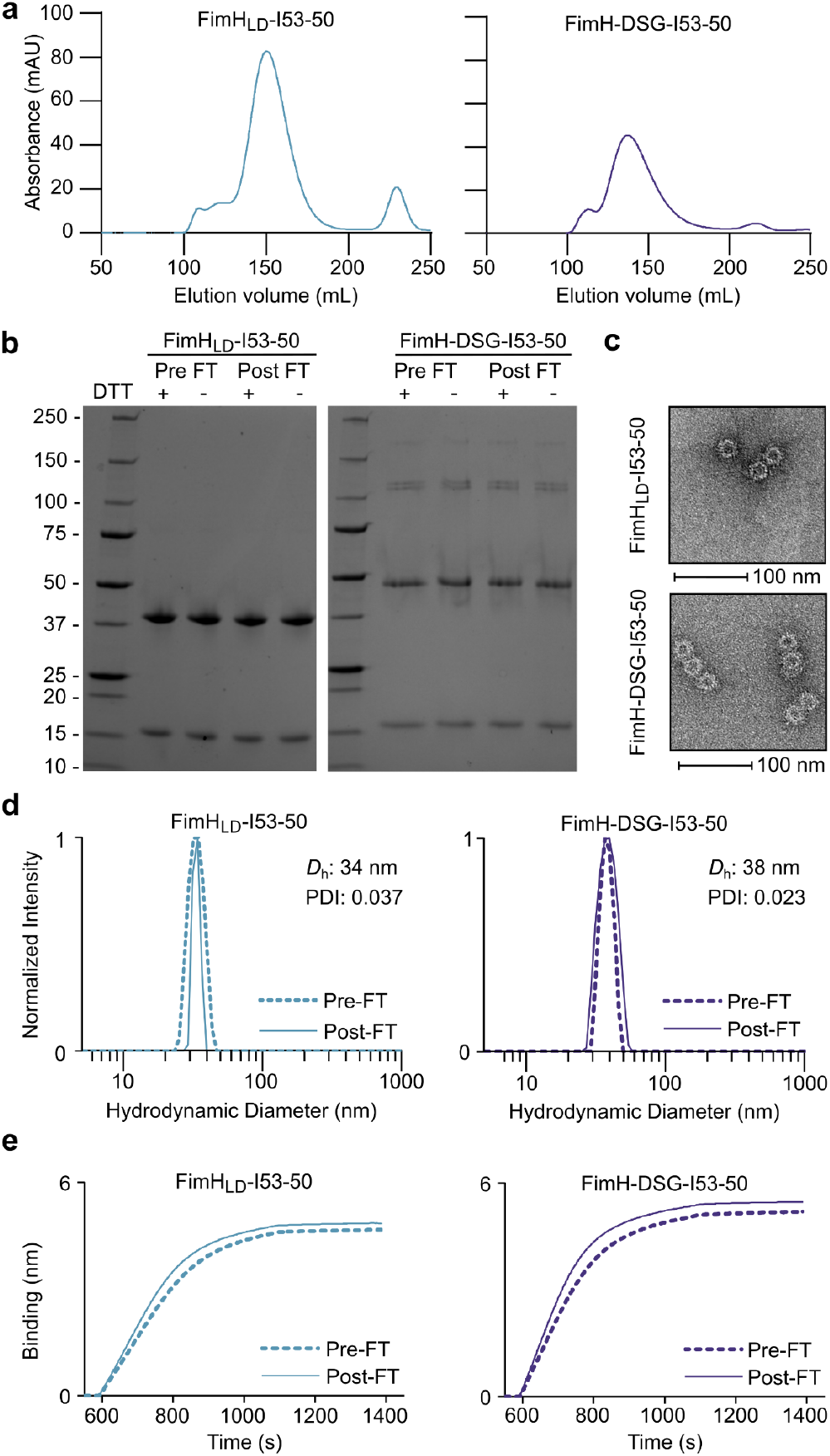
Characterization of FimH_LD_- and FimH-DSG-I53-50 nanoparticles. **a** Preparative SEC of FimH_LD_- and FimH-DSG-I53-50 nanoparticles. **b** SDS-PAGE of pre- and post-freeze/thaw (FT) samples, with (+) and without (-) 10 mM dithiothreitol (DTT). **c** Representative electron micrographs of negatively stained nanoparticles. **d** DLS of both nanoparticle immunogens pre- and post-FT. *D*_h_, hydrodynamic diameter; PDI, polydispersity index. **e** Biolayer interferometry (BLI) of mAb 926 binding to both nanoparticle immunogens pre- and post-FT.

### FimH_LD_-I53-50 and FimH-DSG-I53-50 nanoparticle immunogens induce functional antibody responses in mice

We compared the immunogenicity of the FimH_LD_ and FimH-DSG nanoparticles to soluble (i.e., non-particulate) antigens in CD-1 mice. Two types of non-particulate immunogens were included: monomers and “non-assembling controls” comprising the trimeric antigen-bearing I53-50A components mixed with 2obx-wt, a version of the I53-50B pentamer lacking the interface that drives nanoparticle assembly^30^ (**Fig. 3a** and **Supplementary Table 1**). Most of the immunogens were adsorbed to Alhydrogel (Al(OH)_3_), while the FimH-DSG monomer was administered with a liposomal formulation of 5 µg of QS-21/monophosphoryl lipid A (MPLA) and FimH_LD_ was formulated separately with both adjuvants. Each dose of nanoparticle and the corresponding non-assembling controls comprised 1 µg of FimH antigen (or 1 µg bare I53-50 scaffold), while the monomeric antigens were administered with a 10 µg priming dose followed by two 1 µg (FimH_LD_) or 5 µg (FimH-DSG) booster doses. These comparisons provided a stringent test of the ability of the I53-50 nanoparticle to improve the immunogenicity of FimH_LD_ and FimH-DSG in lieu of higher dosages and stronger adjuvants.

**Fig. 3.**
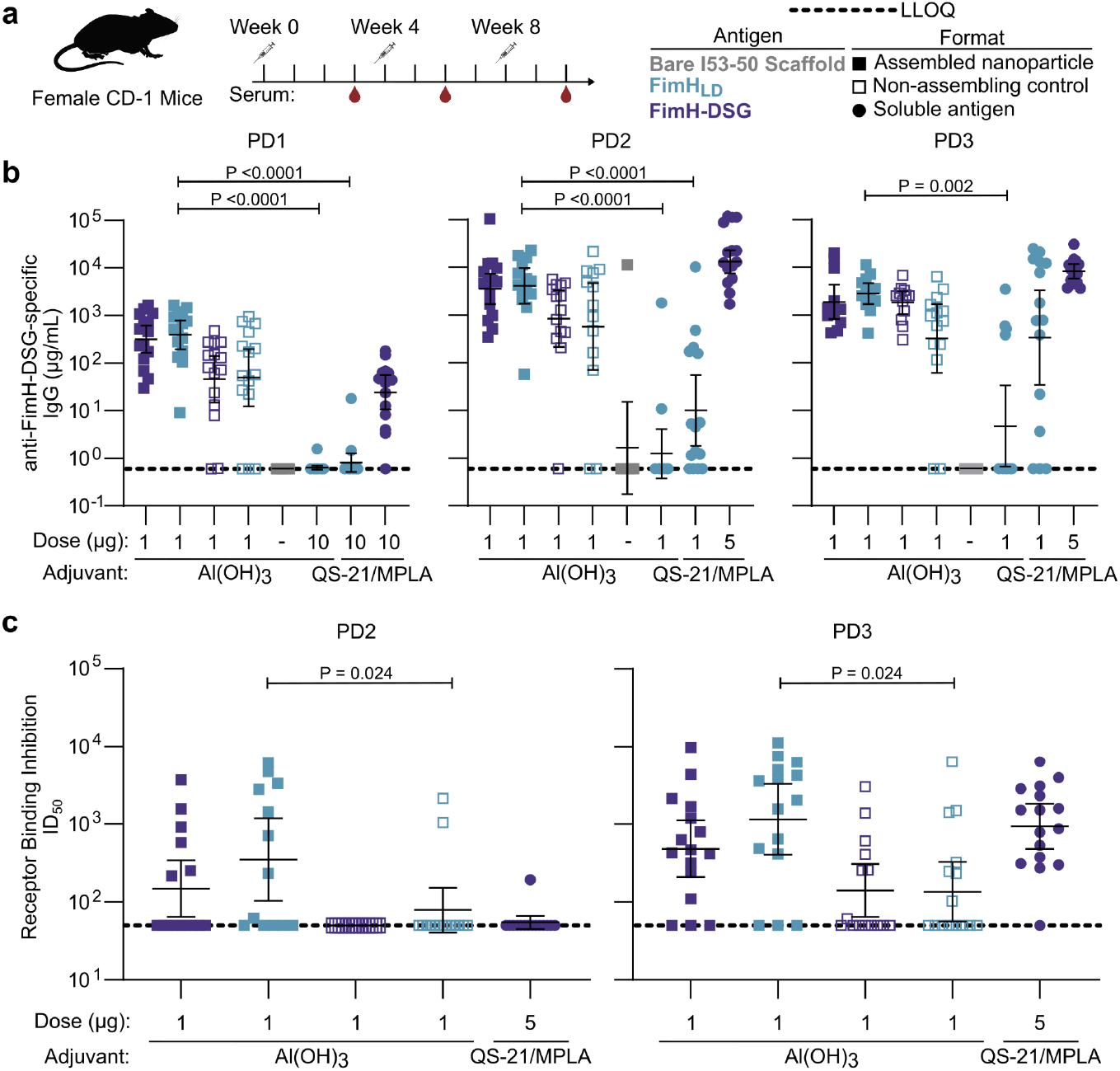
Immunogenicity of FimH_LD_- and FimH-DSG-I53-50 nanoparticles in mice. **a** Study timeline. **b** Post-dose 1 (PD1; Week 3), post-dose 2 (PD2; Week 6), and post-dose 3 (PD3; Week 10) FimH-DSG IgG titers in CD-1 mice, measured by Luminex assay. Each symbol represents an individual animal, and the GMT from each group is indicated by a horizontal line with error bars indicating a confidence interval of 95%. Lower limit of quantification represented with dashed line. **c** Post-dose 2 and post-dose 3 serum receptor binding inhibition titers. Data are presented as in **b**. All groups were compared with a Kruskal-Wallis test followed by Dunn’s multiple-comparisons test. Panel **a** partially created in BioRender^31^.

Groups of 15 female CD-1 mice were immunized intramuscularly at weeks 0, 4, and 8. Three weeks post dose 1, both nanoparticle immunogens elicited serum anti-FimH-DSG IgG responses with geometric mean titers (GMTs) between 1×10^2^ and 1×10^3^ µg/mL (**Fig. 3b**). The non-assembling control immunogens elicited slightly lower titers (GMTs between 1×10^1^ and 1×10^2^ µg/mL). Although both FimH_LD_ monomer formulations yielded little to no response following the first dose, the group receiving 10 µg of monomeric FimH-DSG adjuvanted by QS-21/MPLA had a GMT of 2.5×10^1^ µg/mL. After the second and third doses, anti-FimH-DSG IgG responses from both nanoparticles reached GMTs between 1×10^3^ and 1×10^4^ µg/mL. The levels of antigen-specific IgG induced by the non-assembling control immunogens reached levels similar to the assembled nanoparticles, particularly after the third dose. Among the groups that received monomeric antigens, FimH-DSG adjuvanted with QS-21/MPLA performed the best, eliciting similar levels of anti-FimH-DSG IgG as the nanoparticles post dose 2 and 3. The two FimH_LD_ monomer groups included some mice that seroconverted as well as mice that did not, with QS-21/MPLA driving a higher rate of seroconversion than Alhydrogel.

To measure the ability of antibodies elicited by FimH constructs to inhibit FimH binding to the cognate mannose ligand, a binding inhibition assay^29^ was performed measuring inhibition of *E. coli* bacterial binding to a coated yeast mannan surface in the presence of diluted mouse serum samples. Yeast mannan is a polysaccharide composed of mannose residues, which has been used to model mannosylated surfaces to which FimH might bind. After two doses of each nanoparticle immunogen, half-maximal inhibitory dilutions (ID_50_) in serum samples from individual mice ranged from undetectable to over 1×10^4^, with 50% of the samples exhibiting receptor-blocking activity (**Fig. 3c**). By contrast, only 3 of the 45 mice that received the non-assembling controls or 10 µg of monomeric FimH-DSG had detectable serum inhibitory activity post dose 2. After the third immunization, most of the sera from the FimH-DSG-I53-50, FimH_LD_-I53-50, and QS-21/MPLA monomeric FimH-DSG groups exhibited functional responses, with GMTs of 4.8×10^2^, 1.1×10^3^, and 9.3×10^2^, respectively. Fewer mice in the non-assembling control groups yielded sera with detectable receptor-blocking activity (14 out of 31 mice across both groups). Given the low levels of antigen-specific IgG elicited by the monomeric FimH_LD_ groups (**Fig. 3b**), these groups were not evaluated in the mannan binding inhibition assay. Together, these data show that both the FimH_LD_ and FimH-DSG nanoparticles elicit a similar or greater antibody response compared with their soluble antigen counterparts at a 10-fold lower dose and, in the case of FimH-DSG, using a more basic adjuvant.

### FimH_LD_-I53-50 and FimH-DSG-I53-50 nanoparticle immunogens induce functional antibody responses in non-human primates

To further evaluate potential differences between the FimH_LD_ and FimH-DSG nanoparticle vaccine candidates, both were tested in a non-human primate (NHP) model that more closely resembles the human immune response^32^. NHPs were vaccinated with a dose of each nanoparticle immunogen equivalent to 50 µg of lectin domain at 0, 4, and 14 weeks, again using Alhydrogel as adjuvant (**Fig. 4a**). As a comparator we included monomeric FimH-DSG in QS-21/MPLA at an equivalent dose of lectin domain, as well as a placebo control group that received a solution of sterile saline.

**Fig. 4.**
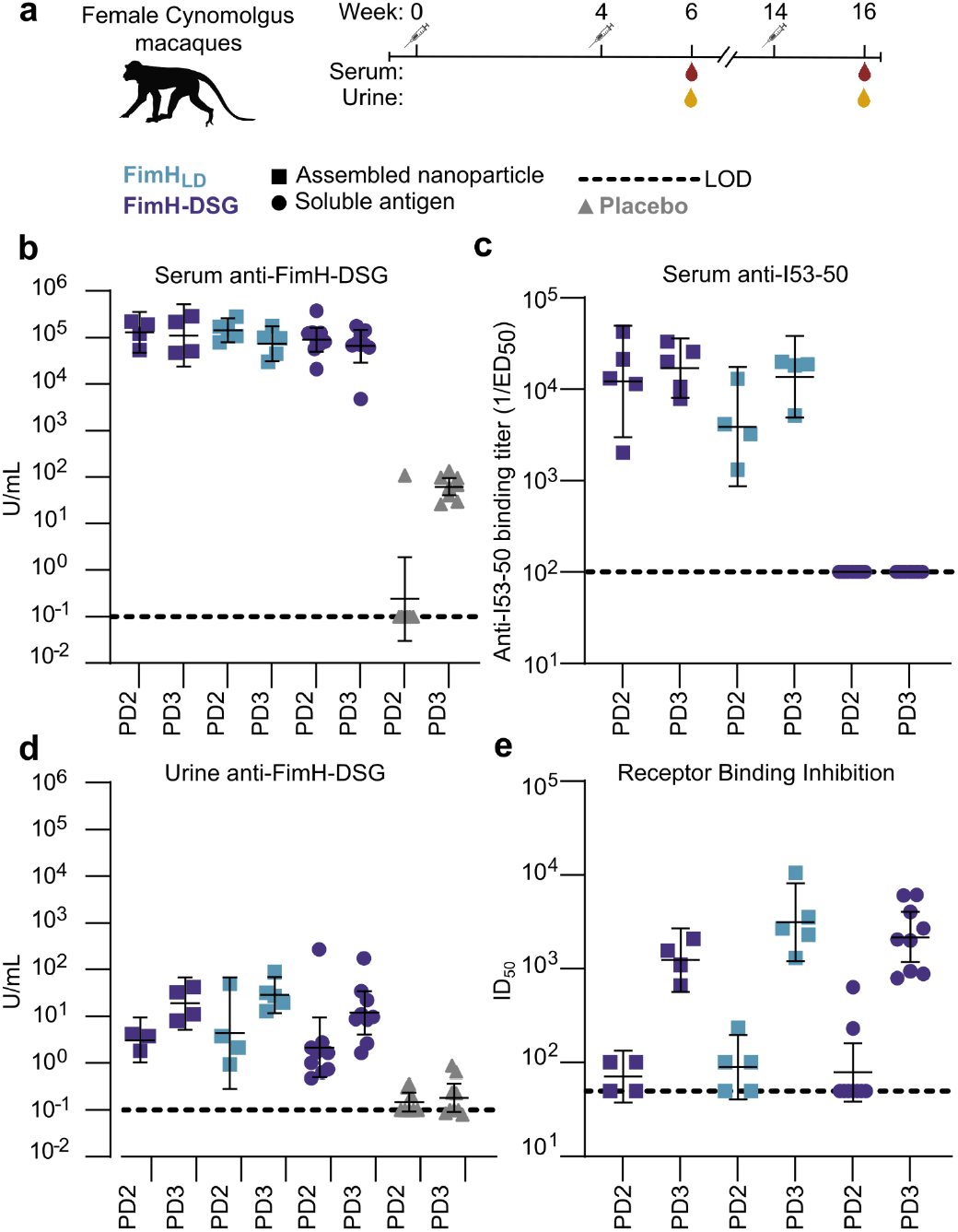
Immunogenicity of FimH-DSG- and FimH_LD_-I53-50 nanoparticles in non-human primates. **a** Study design and timeline. **b** Post-dose 2 (Week 6), and post-dose 3 (Week 16) serum anti-FimH-DSG IgG titers, measured by Luminex assay. Each symbol represents an individual animal, and the GMT from each group is indicated by a horizontal line with error bars indicating a confidence interval of 95%. **c** Post-dose 2 and post-dose 3 serum anti-I53-50 IgG binding titers, measured by ELISA. Data presented as in panel **b. d** Post-dose 2 and post-dose 3 anti-FimH-DSG IgG titers in urine, measured by Luminex assay. Data presented as in panel **b. e** Post-dose 2 and post-dose 3 serum receptor binding inhibition titers. Data presented as in panel **b**. No significant differences between groups were detected by Kruskal-Wallis test followed by Dunn’s multiple-comparisons test. Panel **a** partially created in BioRender^34^.

Post-dose 2 and 3, levels of FimH-DSG-specific IgG in serum were uniformly high (approximately 1×10^5^ U/mL) in NHPs that received FimH immunogens; no significant differences between groups were observed (**Fig. 4b**). We also measured the levels of serum antibody specific to the I53-50 nanoparticle and found that both of the nanoparticle immunogens elicited robust anti-scaffold antibody responses, as we have observed previously^17,33^, while the NHPs receiving the FimH-DSG monomer exhibited no detectable reactivity (**Fig. 4c**). Levels of FimH-DSG-specific IgG in the urine of the vaccinated NHPs were considerably lower overall, but did increase from PD2 to PD3; again no significant differences were observed between groups (**Fig. 4d**). By contrast, urine from NHPs that received placebo contained nearly undetectable levels of antigen-specific IgG.

Measuring serum receptor binding inhibition activity using the yeast mannan binding assay showed that all three immunogens elicited negligible titers after two doses (ID_50_s of roughly 1×10^2^ were sporadically observed) (**Fig. 4e**). A third immunization increased receptor-blocking titers considerably, to between 1×10^3^ and 1×10^4^ for all three groups. Together, these data are similar to what we observed in mice and show that both the FimH_LD_ and FimH-DSG nanoparticles elicit similar levels of functional antibody as an equivalent dose of FimH-DSG in a more potent adjuvant.

## Discussion

We recently reported the design of a conformationally stabilized variant of FimH comprising both the lectin and pilin domains, FimH-DSG (ref. ^12^). Here we compared the immunogenicity of two-component nanoparticle immunogens displaying the stabilized antigen (FimH-DSG) or a traditional antigen comprising only the lectin domain (FimH_LD_) to non-particulate immunogens. Our data showed that alum formulations comprising FimH_LD_ or FimH-DSG displayed on the I53-50 nanoparticle were comparably immunogenic to monomeric FimH-DSG in a more potent adjuvant in mice and NHPs. In mice, an indistinguishable immune response was observed even at a 10-fold lower priming dose of nanoparticle immunogen. These findings are consistent with a previous study in which an alum-adjuvanted nanoparticle immunogen displaying the SARS-CoV-2 RBD elicited higher levels of neutralizing activity than a five-fold higher dose of monomeric RBD formulated on alum + CpG (ref. ^35^). That study and others also found that the responses elicited by RBD nanoparticle vaccines can be further improved through the use of more potent adjuvants than alum^36,37^. However, given the increased formulation complexity and potential reactogenicity that may accompany some adjuvants, the inherent enhanced immunogenicity of nanoparticle immunogens may provide an advantage.

We found that both nanoparticle immunogens elicited high levels of antigen-specific antibody and receptor-blocking activity in mice and NHPs, with no discernible differences in terms of immunogenicity. These data recall similar observations made previously with nanoparticle immunogens displaying the wild-type RBD from the ancestral SARS-CoV-2 virus or a stabilized version containing three repacking mutations in the linoleic acid binding site^30^. In that case, like the present work, the stabilized antigen exhibited advantages in manufacturability and stability, including higher protein yield and increased thermal denaturation, yet these did not translate to improved immunogenicity when displayed on a protein nanoparticle^30^. However, this is not always the case: other studies have found that nanoparticle immunogens displaying stabilized antigens can induce more potent immune responses^38,39^. Nevertheless, the repacking mutations in the previously described SARS-CoV-2 RBD nanoparticle immunogen proved to be critical to the scalable manufacture of a tetravalent pan-sarbecovirus vaccine candidate that recently entered clinical trials^40,41^. Furthermore, the stabilizing mutations were also required for productive secretion of one-component, “mRNA-launched” nanoparticle immunogens displaying the RBD^42^. These examples highlight the potential value of the improved manufacturability of stabilized FimH antigens and nanoparticle immunogens.

We found that in addition to the induction of immune responses to the FimH antigen, immunization with the nanoparticle vaccines also generated antibodies specific to the scaffold. As is documented for glycoconjugate vaccines, antibodies against the carrier protein can either enhance or suppress immune responses to the coupled polysaccharide antigen^43,44^. However, several previous studies with protein nanoparticle immunogens showed that pre-existing immunity against the nanoparticle scaffold did not deleteriously affect antigen-specific responses^17,45^. Furthermore, in clinical trials of the malaria vaccine RTS,S—which displays a portion of the *P. falciparum* CSP protein on a self-assembling hepatitis B surface antigen scaffold^46^—levels of antigen-specific antibodies were either not significantly affected^47–49^ or actually increased^50^ by pre-existing anti-scaffold immunity. Nevertheless, whether anti-scaffold antibodies would complicate sequential immunization with multiple heterologous vaccines utilizing the same nanoparticle scaffold remains an open question.

Multivalent antigen display on protein nanoparticle scaffolds is a widely used approach for enhancing the immunogenicity of subunit vaccines^51^. A theme that has emerged from many of these studies—including the present work—is that small, monomeric antigens tend to benefit the most from nanoparticle display^52^. This is likely due to a combination of antigen receptor clustering and improved trafficking of particulate immunogens to lymph nodes and B cell follicles compared to soluble (i.e., monomeric) antigen^53^. However, the majority of reports to date have focused on displaying viral or parasitic antigens, with few examples of bacterial antigens displayed on commonly used protein nanoparticle scaffolds such as I53-50, ferritin, or Hepatitis B core antigen^54,55^. One such study recently reported that display of the small, monomeric *N. meningitidis* factor H-binding protein (fHbp) on I53-50 elicited bactericidal antibody responses with enhanced magnitude and breadth compared to monomeric and trimeric fHbp immunogens or a licensed MenB vaccine with similar antigenic composition^23^. Our data provide another example of multivalent display increasing the immunogenicity of a small bacterial antigen. Given the wide variety of such antigens that—like FimH—have been shown to be weakly immunogenic and partially protective in soluble form absent a potent adjuvant, these results suggest that nanoparticle scaffolds like I53-50 could be broadly useful for designing antibacterial vaccines that elicit immune responses with improved magnitude, quality, and breadth.

## Acknowledgements

This study was supported by Pfizer Inc.

## Data availability

All data needed to evaluate the conclusions in the paper are present in the paper and/or the Supplementary Materials.

## Methods

### Expression and purification of FimH_LD_- and FimH-DSG-I53-50A trimers

Sequences for FimH_LD_ and FimH-DSG were genetically fused to the N terminus of the trimeric I53-50A nanoparticle component using linkers of 10 glycine and serine residues (**Table S1**). Each I53-50A fusion protein also bore a C-terminal hexa-histidine tag. All sequences were cloned into pcDNA3.1 via Gibson assembly and codon-optimized for mammalian expression. 1 L of ExpiCHO cells were transfected with 0.5 g plasmid DNA encoding FimH_LD_-I53-50A or FimH-DSG-I53-50A according to manufacturer’s instructions. Supernatants were harvested after 8 days, at 80% cell viability, and were clarified with 0.22 µm Sartoclear Dynamics® LabV filters. FimH_LD_- and FimH-DSG-I53-50A were then purified using a two-step purification strategy. The first step was done using an AKTA-Avant FPLC instrument (Cytiva) and a 5 mL HisTrap HP column (Cytiva) (Buffers used: Buffer A: 25 mM Tris pH 8, 300 mM NaCl, 20 mM Imidazole. Buffer B: 25 mM Tris pH 8, 300 mM NaCl, 500 mM Imidazole). Fractions were run on an SDS-PAGE gel, followed by Coomassie blue staining. Desired fractions were then pooled together and underwent a second step of polishing by Hydrophobic Interaction Chromatography (HIC) using a Phenyl Sepharose 5 mL column (Cytiva) on an AKTA-Avant FPLC instrument (Cytiva). (Buffers used: Sample Dilution Buffer: 25 mM Tris pH 8, 600 mM Na-Citrate. Buffer A/Sample buffer: 25 mM Tris pH 8, 300 mM Na-Citrate. Buffer B (Elution Buffer): 25 mM Tris pH 8, 10% v/v glycerol). Fractions were run on an SDS-PAGE gel, followed by Coomassie blue staining. Desired fractions were then pooled together and dialyzed into their final storage buffer (25 mM Tris pH 8, 150 mM NaCl). Samples were further treated with Proteus NoEndo Spin Column Kits (Protein Ark) to remove residual endotoxin. The final FimH_LD_- and FimH-DSG-I53-50A products were ∼95% pure based on SDS-PAGE followed by Coomassie blue staining.

### Production of pentamers

Plasmids encoding 2obx-wt and I53-50B.4PT1 (**Table S1**) were synthesized by GenScript and cloned into pET29b between the NdeI and XhoI restriction sites. I53-50B.4PT1 was incorporated with a double stop codon directly before the C-terminal polyhistidine tag to make a tagless construct. The pentamers were expressed in Lemo21(DE3) cells (NEB) using LB medium (10 g Tryptone, 5 g Yeast Extract, 10 g NaCl per liter) in a 10 L BioFlo 320 Fermenter (Eppendorf). At inoculation, culture conditions were set to 37°C, 225 rpm impeller speed, and 5 SLPM (standard liter per minute) gas flow, with O_2_ supplementation active as part of the dissolved oxygen (DO) cascade. At the onset of a dissolved oxygen spike (OD ∼12), the culture was supplemented with 100 mL of 100% glycerol and induced with 1 mM isopropyl β-D-1-thiogalactopyranoside (IPTG). Simultaneously, the temperature was lowered to 18°C and O_2_ supplementation was stopped. Protein expression continued overnight until reaching an OD ∼20.

2obx-wt cells were harvested and lysed by microfluidization using a Microfluidics M110P at 18,000 psi in 50 mM Tris pH 8, 500 mM NaCl, 30 mM imidazole, 1 mM PMSF, 0.75% w/v CHAPS (3-[(3-Cholamidopropyl)dimethylammonio]-1-propanesulfonate). Lysates were clarified by centrifugation at 24,000 g for 30 min and applied to a 2.6 × 10 cm Ni Sepharose 6 FF column (Cytiva) for purification by IMAC on an AKTA Avant150 FPLC system (Cytiva). Protein of interest was eluted over a linear gradient of 30 mM to 500 mM imidazole in a background of 50 mM Tris pH 8, 500 mM NaCl, 0.75% w/v CHAPS buffer.

I53-50B.4PT1 cells were harvested by centrifugation, and pellets were resuspended in dPBS, homogenized, and lysed using a Microfluidics M110P microfluidizer at 18,000 psi for three passes. Lysates were clarified by centrifugation (24,000 g, 30 min, 4°C), and the supernatant was discarded to isolate inclusion bodies. Inclusion bodies were first washed with dPBS pH 8.0 supplemented with 0.1% Triton X-100, followed by centrifugation for sample clarification. The resulting pellet was then washed with dPBS pH 8.0 supplemented with 1 M NaCl. After the second wash, pentamer protein was extracted from the pellet using dPBS pH 8.0, 2 M urea, 0.75% w/v CHAPS. Extracted protein was loaded onto a DEAE Sepharose FF column (Cytiva) using an AKTA Avant150 FPLC system (Cytiva). After binding, the column was washed with 5 column volumes (CV) of dPBS pH 8.0 + 0.1% Triton X-100, followed by 5 CV of dPBS pH 8.0 + 0.75% w/v CHAPs. Elution was performed using 3 CV of dPBS pH 8.0 + 500 mM NaCl.

Eluted fractions for both 2obx-wt and I53-50B.4PT1 were confirmed by SDS-PAGE, pooled, concentrated using 10K MWCO centrifugal filters (Millipore), sterile-filtered (0.22 μm), and applied to a HiLoad S200 pg GL SEC column (Cytiva) using 50 mM Tris pH 8, 500 mM NaCl, 0.75% w/v CHAPS buffer. After sizing, peak fractions for each protein were confirmed by SDS-PAGE, pooled, concentrated using 10K MWCO centrifugal filters (Millipore) to 50 µM, sterile-filtered (0.22 μm), and assessed for low endotoxin levels prior to mixing (2obx-wt) or assembly (I53-50B.4PT1) with FimH_LD_-I53-50A and FimH-DSG-I53-50A trimers.

### In vitro nanoparticle assembly

Total protein concentration of purified individual nanoparticle components was determined by measuring absorbance at 280 nm using a UV/vis spectrophotometer (Agilent Cary 3500) and calculated extinction coefficients^56^. The assembly steps were performed at room temperature with addition in the following order: FimH_LD_-I53-50A or FimH-DSG-I53-50A trimeric fusion protein, followed by additional assembly buffer as needed to achieve desired final buffer concentration (50 mM Tris pH 7.4, 185 mM NaCl, 100 mM L-arginine, 4.5% v/v glycerol, 0.75% w/v CHAPS), and finally I53-50B.4PT1 pentameric component, with a molar ratio of FimH_LD_-I53-50A or FimH-DSG-I53-50A to I53-50B.4PT1 of 1.1:1. Assemblies were incubated at room temperature for 1 hour before subsequent purification by SEC on a Superose 6 prep grade XK 26/70 column (Cytiva) in order to remove residual unassembled component. Assembled particles elute at ∼140 mL on this column. Pooled fractions were subjected to two rounds of dialysis into their storage buffer of 50 mM Tris pH 8, 150 mM NaCl, 100 mM

L-arginine, 4.5% v/v glycerol. Assembled nanoparticles were sterile filtered (0.22 μm) immediately prior to column application and following pooling and dialysis of fractions.

### Preparation of non-assembling controls

FimH_LD_-I53-50A trimers or FimH-DSG-I53-50A trimers were each combined with an assembly-incompetent wild-type pentamer, 2obx-wt, as described above. Lack of assembly was confirmed by HPLC, where two separate peaks were present for each nanoparticle component.

### Negative stain electron microscopy

FimH_LD_-I53-50 nanoparticles and FimH-DSG-I53-50 nanoparticles were diluted to 0.1 mg/mL in 50 mM Tris pH 8, 150 mM NaCl, 100 mM L-arginine, 4.5% v/v glycerol prior to application of 3 µL of sample onto freshly glow-discharged 300-mesh copper grids. Sample was incubated on the grid for 30 seconds before being blotted away with filter paper (Whatman). Grids were stained three times with 3 μL of 2% w/v uranyl formate stain, blotting immediately with filter paper between each application. The grids were allowed to dry for 1 min. Prepared grids were imaged in a Talos model L120C electron microscope at 57,000× magnification.

### Bio-layer interferometry

mAb 926 was reconstructed from previously described sequences^29,57^. Heavy and light chain sequences were cloned into a pTT5 plasmid with a humanized IgG1 backbone and the antibody was expressed in mammalian cells and purified (LakePharma)^29^. Binding of mAb 926 to FimH_LD_-I53-50 nanoparticles and FimH-DSG-I53-50 nanoparticles was analyzed using bio-layer interferometry with an Octet Red96e System (FortéBio) at ambient temperature with shaking at 1000 rpm. FimH_LD_-I53-50 nanoparticles and FimH-DSG-I53-50 nanoparticles both pre- and post-flash freeze and thaw were each diluted to 100 nM in Kinetics buffer (99.03% HBS-EP+buffer (Cytiva), 0.05% w/v nonfat milk, and 0.02% w/v sodium azide). mAb 926 was diluted to 10 μg/mL in Kinetics buffer. Reagents were applied to a black 96-well Greiner Bio-one microplate at 200 μL per well. Protein A biosensor tips were prewetted in Kinetics buffer for 10 mins. The Protein A tips were loaded with diluted mAb 926 for 500 seconds and washed with Kinetics buffer for 60 seconds. The association step was performed by dipping the Protein A tips with immobilized mAb 926 into diluted nanoparticle for 500 seconds, then dissociation was measured by inserting the tips back into Kinetics buffer for 300 seconds. The data were baseline subtracted and the plots fitted using the Pall Forté Bio/Sartorius analysis software (version 12.0).

### Dynamic light scattering

Dynamic light scattering (DLS) was used to measure hydrodynamic diameter (D_h_) and Polydispersity Index (PDI) of FimH_LD_-I53-50 nanoparticle and FimH-DSG-I53-50 nanoparticle samples on an UNcle Nano-DSF (UNchained Laboratories) at 20°C. Samples were applied to a 8.8 mL quartz capillary cassette (UNi, UNchained Laboratories) and measured with 10 acquisitions of 5 s each, using auto-attenuation of the laser. Increased viscosity due to 4.5% v/v glycerol in the nanoparticle buffer was accounted for by the UNcle Client software in D_h_ measurements.

### Endotoxin measurements of protein nanoparticle vaccine candidates

Endotoxin levels in protein nanoparticle samples were measured using the EndoSafe Nexgen-MCS System (Charles River) prior to immunogen formulation. Samples were diluted 1:100 in Endotoxin-free LAL reagent water, and applied into wells of an EndoSafe LAL reagent cartridge. Charles River EndoScan-V software was used to analyze endotoxin content, automatically back-calculating for the dilution factor. Endotoxin values were reported as EU/mL which were then converted to EU/mg based on UV/vis concentration measurements.

### UV/vis spectroscopy

Ultraviolet-visible spectra were measured using an Agilent Technologies Cary 3500. Samples were applied to a 10 mm, 50 mL quartz cell (Starna Cells, Inc.) and absorbance was measured from 180 to 1000 nm. Net absorbance at 280 nm, obtained from measurement and single reference wavelength baseline subtraction, was used with calculated extinction coefficients and molecular weights to obtain protein concentration. All data produced from the UV/vis instrument was processed in the 845x UV/visible System software.

### Animal ethics statement

Mouse and NHP immunogenicity studies were performed at Pfizer, Pearl River, NY, which is accredited by the Association for Assessment and Accreditation of Laboratory Animal Care (AAALAC). All procedures performed on mice and NHPs were in accordance with local regulations and established guidelines and were reviewed and approved by an Institutional Animal Care and Use Committee (IACUC). The work was in accordance with United States Department of Agriculture Animal Welfare Act and Regulations and the NIH Guidelines for Research Involving Recombinant DNA Molecules, and Biosafety in Microbiological and Biomedical Laboratories.

### Mouse immunization

CD-1 mice (15 per group, female, 8-10 weeks of age) were immunized intramuscularly at 0, 4 and 8 weeks with 1 μg FimH_LD_ antigen equivalent per dose of FimH_LD_ or FimH-DSG nanoparticles or non-assembling controls formulated with aluminum hydroxide, monomeric FimH_LD_ (10 μg prime/1 μg boosts) or monomeric FimH-DSG (10 μg prime/5 μg boosts) formulated with liposomal QS-21/MPLA, or FimH_LD_ (10 μg prime/1 μg boosts) formulated with aluminum hydroxide. Sera were collected two weeks after each dose, and stored at -80°C until use.

### NHP immunization

Cynomolgus macaques (female, 5-9 years of age, 2.8-5.7 kg) were immunized intramuscularly on weeks 0, 4 or 14 with saline placebo (9 per group), 50 μg FimH_LD_ equivalent per dose of FimH_LD_ (5 per group) or FimH-DSG (4 per group) antigen delivered in a nanoparticle presentation adjuvanted with aluminum hydroxide or as monomeric FimH-DSG (9 per group) formulated with liposomal QS-21/MPLA adjuvant. Sera were taken on weeks -2, 0, 3, 4, 6, 8, 13, 14, and 16 via the femoral vein, and stored at -80°C until use.

### Anti-I53-50 scaffold ELISA

For ELISAs against bare I53-50 scaffold, 100 μL of 2 μg/mL I53-50 nanoparticles were plated onto 94-well Nunc-Maxisorp (ThermoFisher) plates in 25 mM Tris pH 8.0, 250 mM NaCl, 5% v/v glycerol and incubated while sealed at room temperature with shaking for 1 hour. After incubation the plates were washed 3× with Tris-Buffered Saline Tween (TBST) using a plate washer (BioTek) and blocked with 5% w/v nonfat dry milk in TBST for 1 hour with shaking at room temperature. Plates were washed 3× with TBST and 1:3 serial dilutions of NHP sera were made in 180 μL TBST starting at 1:100 and incubated for 1 hour. Plates were washed 3× with TBST and anti-human horseradish peroxidase conjugated antibodies (Thermo Fisher) added to each well before incubating at room temperature with shaking for 30 minutes. Plates were washed 3× in TBST and 100 µL TMB (SeraCare) was added to each well for 3 minutes at room temperature. The reaction was quenched with the addition of 100 µL of 1 N HCl. Plates were read immediately at 450 nm on an Epoch plate reader (BioTek) and data plotted and fit in Prism (GraphPad) using nonlinear regression sigmoidal, 4PL, X is log(concentration) to determine ED_50_ values from curve fits.

### Serology

Quantification of serum anti-FimH IgG titers in mice were conducted as follows. Briefly, FimH-DSG was immobilized on Luminex bead microspheres with EDC/NHS. FimH-DSG-coupled beads were incubated with serially diluted individual mouse sera or FimH control mAb with shaking at 4°C for 18 hours. After washing, bound FimH-specific IgG was detected with a PE-conjugated goat anti-mouse IgG mouse secondary antibody. Microplates were read on a FlexMap 3D instrument (BioRad). A FimH-specific mouse IgG mAb was used as an internal standard to quantify anti-FimH IgG levels. NHP IgG titers for FimH in serum and urine were calculated similarly, as previously described^4^. Receptor binding inhibition in sera from mice and NHPs was assessed as previously described^4^.

### Quantification and statistical analysis

Kruskal-Wallis tests followed by Dunn’s multiple-comparisons were performed to compare all groups to determine whether they were statistically different for ELISA and neutralization experiments.

## Supplementary Materials

**Table S1.**
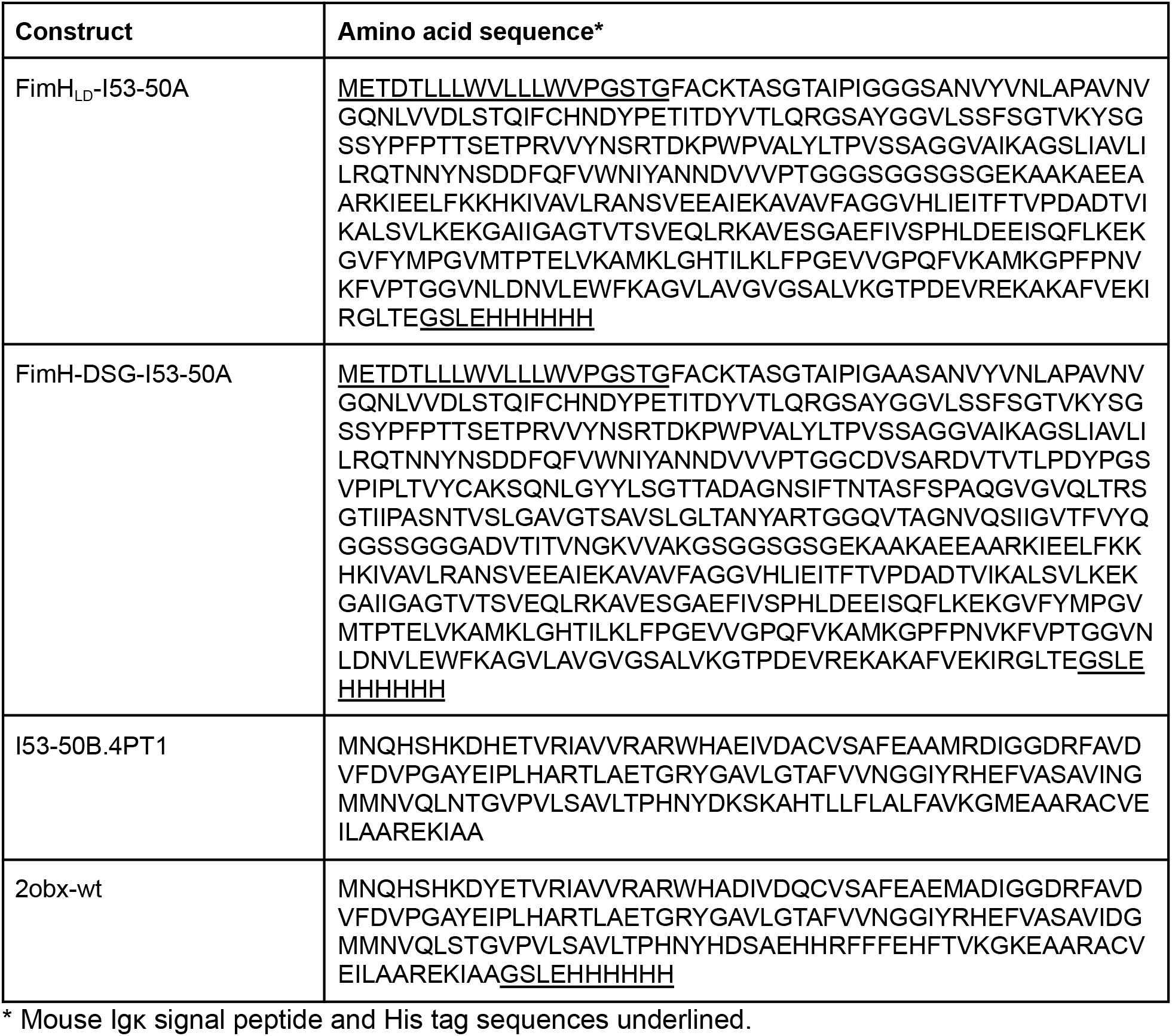
Amino acid sequences of FimH-I53-50A and I53-50B components.

